# Challenges in Improving Genomic Literacy: Results from National and Regional Surveys of Genomic Knowledge, Attitudes, Concerns, and Behaviors

**DOI:** 10.1101/2022.08.26.505444

**Authors:** Joseph Jaeger, Amanda Hellwig, Elizabeth Schiavoni, Bridget Brace-MacDonald, Natalie A. Lamb, Laurene Tumiel-Berhalter, Marc S. Halfon, Arun Vishwanath, Jennifer A. Surtees

**Affiliations:** University at Buffalo’s Genome, Environment and Microbiome (GEM) Community of Excellence, University at Buffalo, Buffalo, NY, USA; Genetics, Genomics and Bioinformatics Graduate Program, Jacobs School of Medicine and Biomedical Sciences, University at Buffalo (SUNY), Buffalo, NY, 14203; Department of Biochemistry, Jacobs School of Medicine and Biomedical Sciences, University at Buffalo, Buffalo, NY, USA; Department of Family Medicine, Jacobs School of Medicine and Biomedical Sciences, University at Buffalo, Buffalo, NY, USA; University at Buffalo’s Clinical and Translational Science Institute, Buffalo, NY, USA; Avant Research Group, LLC, Buffalo, NY, USA

**Keywords:** genomic literacy, positive genomic health behaviors, Western New York, genetic testing

## Abstract

**Purpose:** Information about genomics is increasingly available to mainstream society, with more and more emphasis on using genomic information to make health care decisions. To determine how prepared people are to use this knowledge to make critical health-related decisions, we assessed the public’s level of genomic literacy and whether this knowledge affects their engagement in behaviors related to genomics, such as getting genetic testing.

**Methods:** A survey assessing perceived and actual knowledge, attitudes, concerns, sources of information, and behaviors related to genomics was administered to national and regional samples of participants. A hierarchical linear regression tested whether knowledge and attitudes predicted engagement in behaviors related to genomics.

**Results:** Participants had good basic knowledge of genetics, though they were less familiar with the term “the human genome.” They also displayed positive attitudes towards genomic research, despite expressing many concerns. Both greater knowledge and more positive attitudes significantly and independently predicted greater engagement in genetic testing and other related proactive health behaviors.

**Conclusion:** Knowledge and concerns about genomics impact the public’s ability and willingness to obtain genetic testing and engage in other proactive health behaviors. The public’s genomic literacy could be enhanced by integrating their knowledge (e.g of DNA) with broader concepts (e.g. the human genome and genomics) and how they relate to health. Future research is needed on interventions that do this, to improve the public’s genomic literacy through relationships that build trust

## Introduction

Genomics is the study of the structure and function of genomes, i.e. all the DNA (genes) encoded by a person’s cells. This includes the ways in which genes interact and influence each other and a person’s environment. Variations in one’s genome may have implications for variations in one’s health. The rapid expansion of innovative research in genomics has vast implications for health care, especially the promise of precision medicine [1, 2]. More than ever, it is vital that the public is genomically literate—that is, they have a “working knowledge of genomic science and its role in society, including personal decision-making” [3]. Indeed, we now live in an information age in which the public has access to an enormous amount of information, including personal health information and results from direct-to-consumer genetic testing, which has shifted the burden of making health-related decisions from health care providers to the individual [4–7]. Although access to genomic information is widespread, the integration of genomics into clinical practice remains challenging [8, 6, 7]. Several studies have demonstrated that health care providers lack the knowledge and/or confidence to incorporate genetics/genomics into their practice [9, 8, 10–17]. Thus, patients must serve as self-advocates and be able to initiate genomic-based conversations with their health care providers.

The ease of access to information about broad and personal genomics may lead us to believe that the public is well-informed on the topic, but previous research showed that this may not be the case. Of the adults surveyed in the United Kingdom that were familiar with the term “genetically modified,” only one-third admitted a good or very good understanding of it [18]. Similarly, American adults displayed a conversational familiarity with terms such as “genetic,” but actual understanding of and ability to define basic genetic concepts was lacking [19], Lea and colleagues [20] similarly concluded that while the public correctly believes that genes can play a causal role in the development of complex diseases, they lack knowledge about basic genetic facts. In fact, a direct comparison showed that individuals perform better on disease-related genomic questions than on more “basic” questions about genes, chromosomes, and cells [21]. The level of genetic numeracy, defined as the ability to understand and estimate genetic risk, may also be low. A survey of university students found that only a small proportion had high genetic numeracy, but they were more willing to receive prenatal testing in the future, indicating the translation of genomic knowledge to behavior [22]. Consistent with this possibility, a recent study found that higher numeracy correlated with correct interpretation of a direct-to-consumer pharmacogenetics test report [23].

The National Human Genome Research Institute defined genomic *health* literacy, which includes genetic numeracy, as the “capacity to obtain, process, understand, and use genomic information for health-related decision making” [3] and made it a research priority, acknowledging that inadequate genomic literacy may limit the positive impact of genomics on health care [24, 3]. The goal of such research is to promote public engagement in genetic testing, including genomic sequencing. More than 85% of those surveyed in the United Kingdom were aware of genetic tests that can predict risk of diseases [18], whereas a much smaller proportion (57%) of a US sample demonstrated awareness of genetic tests [25]. However, those surveyed in the United States were more aware of genetic tests to determine disease risk than of tests to determine treatment and drug efficacy, an aspect of precision medicine [25]. Although the public is generally positive towards the use of genetic testing and the objectives of genetic research [21], a variety of concerns have been expressed regarding confidentiality and privacy, trust, not knowing enough about the research being performed, the quality of genetic tests, and the possible use of genetic information as discrimination for employment or obtaining health insurance [1, 26, 21, 18]. However, previous studies had relatively small sample sizes, making it challenging to generalize the results.

To assess the public’s knowledge, attitudes, concerns, sources of information, and behavior related to genomics, we surveyed national (United States) and regional (Erie County in Western New York) populations. We compared the local findings to the national baseline and explored how knowledge of and attitudes towards genomics influence whether individuals act on that knowledge through positive genomic health behaviors. These data will also serve as a baseline against which to measure the local outreach efforts of the Genome, Environment and Microbiome Community of Excellence at the University at Buffalo. Our mission is to advance genome and microbiome research in Buffalo and beyond and to promote genome and microbiome literacy, ensuring the equitable distribution of these scientific advances throughout society.

## Methods

A telephone survey company (Portable Insights) was used to recruit a random sample from US landline telephone numbers. Participants were recruited for both a national (United States) sample and a separate regional (Erie County, NY) sample. Participants were contacted via telephone, and informed consent was obtained for participation (UB IRB Study #00000605). The surveys were conducted in the spring of 2017.

### Measures

Participants were asked to provide demographic information, information about their personal health such as having a genetic disorder and having a gene related to a disease, and responses to the following measures:

#### Perceived knowledge

Participants were asked to rate their understanding of four terms associated with genomic science: human DNA, genetic testing, genetically modified organisms (GMOs), and the human genome (Table S1). Ratings were on a scale of 1–5, with 1 corresponding to “have not heard the term” and 5 indicating “excellent understanding.” These items were summed to provide a single perceived knowledge score for each individual, with total scores ranging from 4 to 20, with a higher value indicating a higher perceived knowledge. The reliability coefficient of this scale was 0.83. Reliability coefficients throughout were determined using Cronbach’s alpha in SPSS.

#### Actual knowledge

Eleven true/false items were used to measure participants’ objective knowledge of genomic science (Table S2). Some items were adapted from previous studies of genomic knowledge [27, 28], whereas others are unique to this study. The items encompassed general genetic concepts, such as “a gene is a piece of DNA” and “a person can have a gene for a specific disease and still be completely healthy.” The sum of correct answers was calculated for each participant to create a total knowledge score, ranging from 0 to 11, with a higher score indicating greater actual knowledge. The reliability coefficient of this scale was 0.38. This may be related to the fact that the questions were all true/false, which provides less variability in the data than those with a scaled response. It is also likely that some of the items in the knowledge scale were measuring a construct separate from genomic knowledge, thus detracting from the internal consistency of the measure. Principal component analysis was performed using the prcomp() function and plotted using the “ggfortify” package in RStudio (Fig. S1).

#### Attitudes

Two items were used to measure participants’ attitudes towards genomics (Table S3). Participants were asked to what extent they agreed that genomics would improve human health, with answers rated on a scale of 1–5, with 1 for “strongly disagree” and 5 for “strongly agree.” Participants were also asked how optimistic they were about the possibility of improved health care as a result of genetic research, with answers again rated on a scale of 1–5, with 1 for “not at all optimistic” and 5 indicating “extremely optimistic.” These items were based partially on instruments used in previous studies of attitudes towards genomics [29, 30]. The sum of the two attitude items was calculated for each participant to create an overall attitude score, ranging from 2 to 10, with a higher value indicating a more positive attitude. The reliability coefficient of this scale was 0.58.

#### Concerns

Participants were asked about their concerns regarding genetic research (Table S4) and the use of their genetic information (Table S5). Several concerns found in previous research were presented to the participants [18], and they were asked to indicate whether each was a concern for them. Participants were allowed to endorse multiple concerns and to provide additional concerns not listed. Some of the additional concerns were similar to those included as options, e.g. concerns about use of genetic information, and were coded as such. The remaining additional concerns were hand-coded separately and are listed as “Unprompted” in Tables S4 and S5. These “unprompted” ideas included concerns about bigotry, discrimination, lack of trust in authorities, regulation, lack of government support, nonconsensual human experimentation, creation of new life (e.g., cloning), genetically modified organisms, and conflict with personal religious beliefs.

#### Sources

Participants were asked two questions to identify their sources of genomic information related to (i) medical research and (ii) the human genome and/or genetic research. Participants indicated all applicable sources: options encompassed print media, digital media, scientific journals, professionals, personal experience, and family and friends (Table S6).

#### Behaviors

Eight behaviors associated with proactive use of genomic knowledge toward health care were presented to participants, and they were asked to rate the degree to which they had considered or actually engaged in these behaviors, from 1 indicting “no, never considered” to 4 for “yes, already done it” (Table S7). Behaviors included getting a genetic test, discussing testing with physicians, friends, or family, participating in research, and making positive lifestyle choices based on family history of disease. Two items, “avoiding recommended vaccinations” and “reading labels to check if a food is genetically modified,” were not considered linked to the other behaviors involving genomic knowledge and were found to detract from the scale’s internal consistency; thus, they were not included in the final behavior scale. The sum of the other six behavior items was calculated for each participant to create a total behavior score ranging from 0 (for declining to answer any question) to 24, with a higher score indicating more positive behaviors. The reliability coefficient of this scale was 0.76.

### Statistical analyses

Anonymous participant data were analyzed using IBM SPSS statistical software. National and regional data were analyzed separately. Because the age distributions of both samples were skewed toward older individuals, the datasets were weighted by age using national and local census data [31]. Response frequencies were obtained for demographic and health-related items, and means and standard deviations were calculated for scale scores (knowledge, attitudes, and behaviors); *t* tests were used to determine whether differences between the two samples’ means were significant. Zero-order correlations were calculated to determine associations between relevant demographics, health variables, knowledge, attitudes, and behavior. Demographic variables that were significantly correlated with behavior were included in the regression.

A hierarchical linear regression investigated whether knowledge and attitudes predicted positive genomic health behavior independent from the effects of demographic characteristics in the regional weighted and national unweighted datasets. Two ordinary least squares regression models were estimated to examine whether the addition of attitudes and knowledge significantly improved the fit of the model and to identify specific variables that uniquely predicted behavior. Regression model 1 estimated the effects of demographic characteristics on behavior, and regression model 2 estimated the effects of knowledge and attitudes on behavior, controlling for demographic characteristics. Ordinal indicators of education (1, less than high school; 2, high school graduate or equivalent; 3, associate’s degree or trade; bachelor’s degree; 4, graduate or professional degree) and annual income (1, <$25,000; 2, $25,000–$49,000; 3, $50,000–$74,999; 4, $75,000–$99,999; 5, ≥$100,000) were included in model 1 along with a dichotomous indicator of gender (1 or 0 for male or female, respectively). A *p* value of <0.05 was considered significant for all tests.

## Results

### Sample characteristics

A total of 1,504 participants completed the survey as part of the national sample. A separate regional sample of 1,000 residents of Erie County, NY, also completed the survey. These two samples were mutually exclusive. Unweighted frequencies for demographic variables of participants in both sample groups are presented in Table 1. Both samples had approximately equal proportions of male and female participants, and White participants were the majority in both samples. A higher percentage of participants in the national sample than those in the regional sample attained at least a bachelor’s degree, and more participants in the national sample reported being in the highest income group. Notably, both samples had greater participation from older individuals. The majority (~80%) of participants in both the national and regional samples reported that their health was good, very good, or excellent. Approximately 15% of participants in both the national and regional samples indicated that they had a genetic disorder and that they had a gene that is related to a disease.

**Table 1.** Characteristics of the samples

| Characteristic | *n* (%)* |  |
| --- | --- | --- |
|  | National dataset<br>(*N* = 1,504) | Regional dataset<br>(*N* = 1,000) |
| Age (years) |  |  |
| 18–24 | 27 (1.8) | 52 (5.2) |
| 25–34 | 35 (2.3) | 170 (4.2) |
| 35–44 | 94 (6.3) | 70 (7.0) |
| 45–54 | 175 (11.6) | 168 (16.8) |
| 55–64 | 358 (23.8) | 251 (25.1) |
| 65–74 | 466 (31.0) | 255 (25.5) |
| ≥75 | 315 (20.9) | 142 (14.2) |
| No answer | 34 (2.3) | 20 (2.0) |
| Sex |  |  |
| Male | 737 (49.0) | 510 (51.0) |
| Female | 767 (51.0) | 490 (49.0) |
| Education |  |  |
| Less than high school | 39 (2.6) | 38 (3.8) |
| High school graduate or equivalent | 426 (28.4) | 392 (39.4) |
| Associate's or vocational degree | 162 (10.8) | 148 (14.9) |
| Bachelor's degree | 410 (27.4) | 215 (21.6) |
| Graduate or professional degree | 461 (30.8) | 202 (20.3) |
| Race |  |  |
| White | 1214 (80.7) | 866 (86.6) |
| Hispanic/Latino | 36 (2.4) | 4 (0.4) |
| Black or African American | 101 (6.7) | 57 (5.7) |
| American Indian or Alaskan Native | 13 (0.9) | 3 (0.3) |
| Asian | 14 (0.9) | 3 (0.3) |
| Native Hawaiian or Pacific Islander | 3 (0.2) | 0 (0.0) |
| Other | 33 (2.2) | 21 (2.1) |
| Multiple races indicated | 46 (3.1) | 23 (2.3) |
| No answer | 44 (2.9) | 23 (2.3) |
| Income |  |  |
| <\$25,000 | 131 (8.7) | 121 (12.1) |
| \$25,000 to \$49,999 | 253 (16.8) | 195 (19.5) |
| \$50,000 to \$74,999 | 236 (15.7) | 173 (17.3) |
| \$75,000 to \$99,999 | 158 (10.5) | 110 (11.0) |
| ≥\$100,000 | 346 (23.0) | 152 (15.2) |
| No answer | 380 (25.3) | 249 (24.9) |
| Quality of Health |  |  |
| Poor | 47 (3.2) | 37 (3.7) |
| Fair | 241 (16.0) | 157 (15.7) |
| Good | 511 (34.0) | 368 (36.8) |
| Very good | 494 (32.8) | 326 (32.6) |
| Excellent | 202 (13.4) | 109 (10.9) |
| No answer | 9 (0.6) | 3 (0.3) |
| Do you have a genetic disorder? |  |  |
| Yes | 227 (15.1) | 154 (15.4) |
| No | 961 (63.9) | 681 (68.1) |
| No answer | 316 (21.0) | 165 (16.5) |
| Do you have a gene that is related to a disease? |  |  |
| Yes | 227 (15.1) | 148 (14.8) |
| No | 737 (49.0) | 573 (57.3) |
| No answer | 540 (35.9) | 279 (27.9) |

\*Unweighted values.

### Perceived knowledge

Participants were asked to rate their understanding of four genomic terms to assess the degree to which they believed they were knowledgeable about these concepts. Four participants in the national sample did not report responses here and were omitted. Participants in the national sample were most familiar with “human DNA,” with >70% responding that they had good (29.2% [438/1,500]) or excellent (44.5% [667/1,500]) understanding of the term (Table S1). Nearly two-thirds of the participants in the national sample reported good (29.7% [445/1,501]) or excellent (33.2% [498/1,501]) understanding of the term “genetic testing.” More than half of national participants reported good (28.9% [434/1,500]) or excellent (26.1% [392/1,500]) understanding of GMOs. Participants felt the least confident in their knowledge of the human genome, with less than half reporting good (23.2% [348/1,500]) or excellent (18.7% [280/1,500]) understanding, and 16.6% (249/1,500) reporting that they had not heard the term at all. Similar patterns were observed in the regional dataset (see Table S1).

### Actual knowledge

A brief true/false quiz assessed participants’ objective knowledge of genomic concepts. On average, participants answered roughly 9 of the 11 items correctly (Table 2, Actual knowledge). This was higher than hypothesized according to previous studies [32, 18]. The statements “A gene is a disease,” “Genes control hereditary characteristics,” “All plants and animals have DNA,” and “A person can have a gene for a disease but be healthy” were most frequently answered correctly (≥93%). Between 70% and 80% of participants correctly identified statements relating genes to DNA (“A gene is a piece of DNA”), relating DNA to the genome, and about the organization of genes in the body (“Different body parts have the same genes”) (Table S2). Only 55.9% (840/1,504) of the national sample and 50.7% (506/1,000) of the regional sample correctly identified “Genes can be altered by the environment” (Table S2). Regional participants scored significantly lower than national participants on the overall measure of actual knowledge (*p* = 0.001; Table 2).

**Table 2.** Weighted mean scale scores

| Category | Weighted $\bar{X}$ (SD) score | | *t* value | *p* value |
| --- | --- | --- | --- | --- |
|  | National dataset | Regional dataset |  |  |
| Actual knowledge | 9.24 (1.40) | 8.98 (1.45) | 4.54 | 0.001 |
| Attitudes | 8.44 (1.41) | 8.25 (1.54) | 3.12 | 0.001 |
| Behavior | 12.70 (5.02) | 11.53 (4.86) | 5.77 | <0.001 |

### Attitudes

The responses to two questions were used to assess participants’ optimism about genomic research. Roughly 90% of the participants in the national sample indicated that they agreed (35.6% [534/1,502]) or strongly agreed (55.3% [830/1,502]) that genomic research would improve human health (Table S3). Additionally, the majority of these participants indicated that they were either optimistic (47.3% [710/1,501]) or extremely optimistic (33.1% [497/1,501]) about the possibility of improved health care as a result of genetic research. Overall, participants had positive attitudes toward genetic research (Table 2). Participants in the regional sample had similar attitudes, but their scores were significantly lower (*p* = 0.01; Table 2).

### Concerns

Participants were asked to state their concerns about genetic research and the use of their genetic information; they could endorse concerns provided to them or state their own. In the national sample, 39.9% (600/1,504) of the participants reported that they did not have any concerns regarding genetic research (Table S4). The most frequently reported concern (15.9% [239/1,504] of participants in the national sample) was a lack of trust in authorities in government, medicine, and/or pharmaceutical companies (unprompted concerns), followed by the concern that such research would be used to create “designer babies” (14.6% [220/1,504] of participants)(prompted concern). The regional sample shared national concerns regarding a lack of trust (9.5% [95/1,000]) and “designer babies” (15.3% [153/1,000]) and also reported concerns with confidentiality (7.9% [79/1,000]) and that genetic research may affect their health insurance premiums, both unprompted concerns (9.9% [99/1,000]) (compared to 9.1% [137/1,504] and 7.6% [114/1,504] of the national participants, respectively) (Table S4).

A similar pattern was observed with regard to concerns about the use of personal genetic material. Although most participants in the national sample reported no concerns (40.8% [613/1,504]), the most frequent concern was related to what would happen to their samples after testing had been completed (18.0% [271/1,504]) (prompted concern; Table S5). Furthermore, participants indicated that they did not trust the authorities who would be handling their samples (14.3% [215/1,504]; both unprompted concerns) or they did not believe that testing and the use of their material would be properly regulated (13.1% [197/1,504]). The participants in the regional sample were similarly most concerned with what happens to their samples after testing (19.5% [195/1,000]) and did not trust authorities to handle their samples properly (13.3% [133/1,000]) (Table S5).

### Sources of information

Several survey questions explored the sources from which participants obtained information on medical and genetic research (Table S6). Among participants in the national sample, the internet was the most frequently reported source of information for medical research (52.7% [792/1,504]), followed closely by health professionals (51.9% [780/1,504]), such as doctors or nurses. Among participants in the regional sample, health professionals were the most frequently reported source (45.7% [457/1,000]), followed by television (44.3% [443/1,000]) and the internet (43.1% [431/1,000]). Regarding the human genome and genomic research, participants in the national sample again reported the internet (45.5% [684/1,504]) as their preferred source of information, followed by magazine articles (38.8% [583/1,504]) and television (38.4% [577/1,504]). Participants in the regional sample similarly reported the internet (37.3% 373/1,000]) as their most popular source for information on genetic research, followed by television (34.4% [344/1,000]) and newspaper articles (33.3% [333/1,000]) (Table S6).

### Positive genomic health behaviors

Participants were asked whether they intended to or had already engaged in eight positive genomic health behaviors; six of these behaviors were included in the final scale. Three participants in the national sample did not respond to this section. Roughly half of the participants in the national sample had never considered getting a genetic test (52.6% [790/1,501]) or seeking information about genetic testing (49.6% [744/1,501]), and most had never considered speaking to their physician about genetic testing (67.6% [1,015/1,501]) (Table S7). The participants were far more likely to speak to family and friends about genetic testing (38.5% [579/1,501]) than to seek out the information on their own (18.9% [284/1,501]). It is worth noting that, because ‘genetic testing’ is self-reported, it is possible that a higher percentage of respondents have received some sort of genetic test but didn’t perceive it as such. Although a majority of participants in the national sample (69.3% [1,041/1,501) had never considered participating in genetic research, 56.3% (845/1,501) indicated that they were actively modifying their lifestyles on the basis of a family history of a certain disease (28.2% [423/1,501] had never considered doing so).

The participants in the regional sample were less likely than the national participants to engage in genomic health behaviors, with 65.6% (656/1,000) reporting that they had never considered getting a genetic test and 63.0% (630/1,000) reporting that they had never considered seeking more information. Nevertheless, 56.2% (562/1,000) reported that they had modified their lifestyle on the basis of a family history of a certain disease (Table S7). As with the national sample, people were more likely to speak with family members (29.5%) than with their physician (14.2%) about genetic testing. Overall, the participants in the national sample were more likely than those in the regional sample to engage in positive genomic health behaviors (*p* > 0.001; Table 2). The total behavior score most strongly correlated with the item “Seeking more information about genetic testing” (*r* = .77) in both samples.

When asked whether they checked to see if foods were genetically modified, 50.4% (757/1,501) of the participants in the national sample reported that they were already engaging in this behavior, whereas 41.3% (620/1,501) had never considered it (Table S7). Most of the national participants (75.8% [1,138/1,501]) reported that they had never considered avoiding vaccinations, whereas 16.0% [241/1,501] indicated that they already had avoided certain vaccinations. It is worth noting that these data were collected prior to the COVID-19 pandemic and attitudes about vaccination may have changed.

### Zero-order correlations

Correlations among demographics, health variables, and total scores of knowledge, attitudes, and behaviors—variables to be included in the regression—were examined using the age-weighted national dataset. Attitudes moderately correlated with knowledge and behaviors, and knowledge and behaviors were also moderately correlated (*r* values between 0.015 and 0.21) (Table 3). Education significantly positively correlated with knowledge, attitudes, and behavior, whereas age negatively correlated with attitudes and behavior. Participants’ self-reports of the quality of their health was slightly negatively correlated with behavior, with healthier individuals less likely to endorse behaviors. Having a genetic disorder or a gene related to a disease was positively correlated with behavior (Table 3).

**Table 3.**
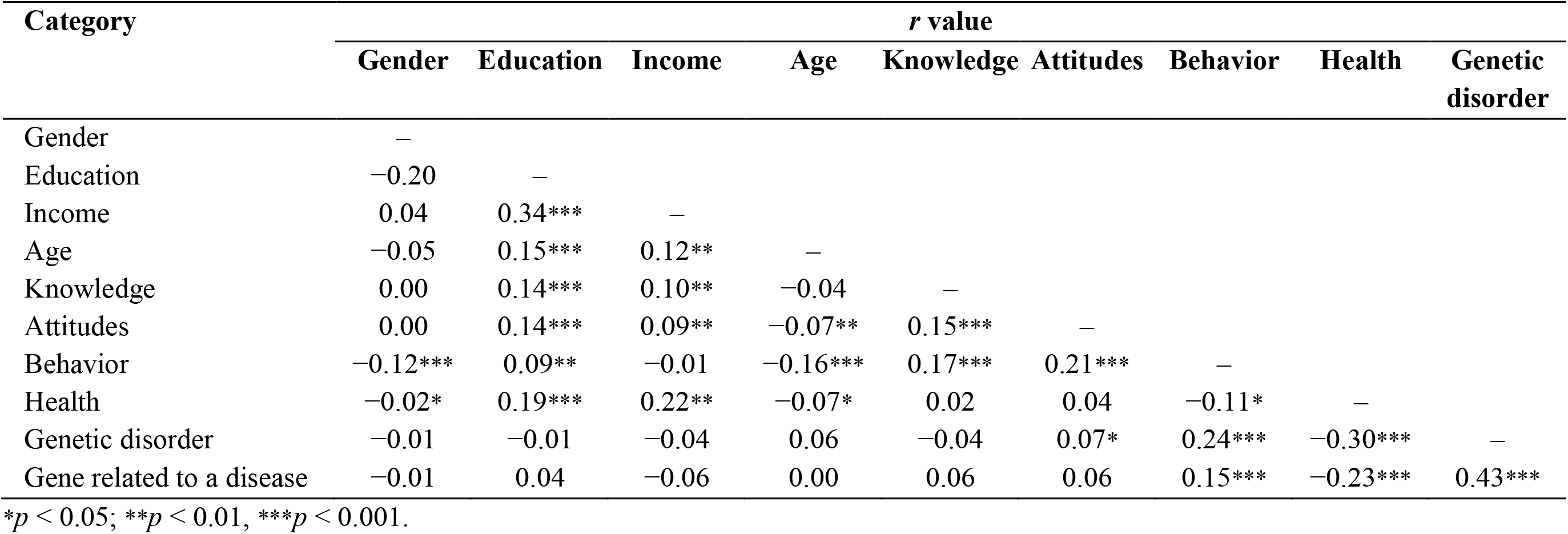
Correlations among demographics, knowledge, attitudes, and behavior in a national dataset

| Category | *r* value |  |  |  |  |  |  |  |  |
| --- | --- | --- | --- | --- | --- | --- | --- | --- | --- |
|  | Gender | Education | Income | Age | Knowledge | Attitudes | Behavior | Health | Genetic disorder |
| Gender | — |  |  |  |  |  |  |  |  |
| Education | −0.20 | — |  |  |  |  |  |  |  |
| Income | 0.04 | 0.34*** | — |  |  |  |  |  |  |
| Age | −0.05 | 0.15*** | 0.12** | — |  |  |  |  |  |
| Knowledge | 0.00 | 0.14*** | 0.10** | −0.04 | — |  |  |  |  |
| Attitudes | 0.00 | 0.14*** | 0.09** | −0.07** | 0.15*** | — |  |  |  |
| Behavior | −0.12*** | 0.09** | −0.01 | −0.16*** | 0.17*** | 0.21*** | — |  |  |
| Health | −0.02* | 0.19*** | 0.22** | −0.07* | 0.02 | 0.04 | −0.11* | — |  |
| Genetic disorder | −0.01 | −0.01 | −0.04 | 0.06 | −0.04 | 0.07* | 0.24*** | −0.30*** | — |
| Gene related to a disease | −0.01 | 0.04 | −0.06 | 0.00 | 0.06 | 0.06 | 0.15*** | −0.23*** | 0.43*** |

\* $p < 0.05$ ; \*\* $p < 0.01$ , \*\*\* $p < 0.001$ .

### Hierarchical linear regression

A hierarchical linear regression examined the effects of knowledge and attitudes on behavior, controlling for demographic factors. The first block of the hierarchical linear regression using the age-weighted national dataset (model 1) determined the effects of demographic variables on behavior. In this first model, there was a significant effect of gender, with males less likely to engage in these behaviors than females (Table 4). There was a significant negative effect of age as well, with older individuals less likely to engage in positive genomic behaviors (Table S7). Level of education predicted greater engagement in the behaviors, with individuals that achieved higher levels of education more likely to engage in positive genomic behaviors. Model 1 explained roughly 6% of the variance in behavior (*R*^2^ = 0.056, *p* < 0.001).

**Table 4.** Results of hierarchical linear regression using national dataset

| Category | $\beta$ value | |
| --- | --- | --- |
|  | Model 1 | Model 2 |
| Gender | −1.316*** | −1.31e2*** |
| Education | 0.473*** | 0.282** |
| Age | −0.497*** | −0.430*** |
| Actual knowledge |  | 0.476*** |
| Attitudes |  | 0.600*** |

\*\* $p < 0.01$ , \*\*\* $p < 0.001$ .

The second block of the hierarchical linear regression (model 2) determined the effects of actual knowledge and attitudes on behavior. After controlling for demographic characteristics, both knowledge and attitudes independently significantly predicted greater engagement in the behaviors (Table 4). Individuals with greater knowledge of basic genetic and genomic concepts were more likely to engage in positive genomic health behaviors, even after controlling for the effects of gender, age, and education. Likewise, individuals with more positive attitudes towards genomic science were more likely to engage in these behaviors after controlling for demographics. The addition of both actual knowledge and attitudes to the model led to a significant increase in variance (5%) accounted for by the model (change in *R*^2^ = .050, *p* < 0.001), with model 2 explaining roughly 11% of the variance in behavior (*R*^2^ = 0.106).

In a supplementary analysis, a multiple regression model was examined (Table 5). This model was identical to model 2 but incorporated the participants’ reports about their health, whether they had a genetic disorder, and whether they had a gene related to a disease. These items had high percentages of missing data and were therefore omitted from the previous regression analysis. This revised model was fitted using only cases with complete data (*N* = 864, 57% of the original national sample size). Having a genetic disorder predicted greater engagement in positive genomic health behavior, controlling for demographic characteristics, knowledge, and attitudes (Table 5). The revised model accounted for 15% of the total variance in behavior (*R*^2^ = 0.153, *p* < .001).

**Table 5.** Results of multiple regression with health questions using national dataset subset

| Category | $\beta$ value |
| --- | --- |
| Gender | −0.985** |
| Education | 0.403** |
| Age | −0.438*** |
| Quality of health | −0.649*** |
| Genetic disorder | 2.842*** |
| Gene related to a disease | −0.144 |
| Actual knowledge | 0.595*** |
| Attitudes | 0.356** |

\*\* $p < 0.01$ , \*\*\* $p < 0.001$ .

The analyses described above were repeated using the regional dataset. The correlation matrix (Table S8), hierarchical regression results (Table S9), and multiple regression results including health questions (Table S10) are available in the supplementary materials. Overall, similar results were observed in both datasets. Notable differences occurred in the multiple regression analyses, which included data for the health questions (Table S10). In the national sample, genomic knowledge remained a significant predictor of behavior when questions about health were included (Table 5). This was not the case in the regional dataset. Likewise, age and self-reported health did not predict behavior in the regional dataset, whereas they did in the national dataset (compare Tables S10 and 5). In the regional dataset, having a gene related to a disease predicted behavior (β = 1.843, *p* < 0.001). The model accounted for 14% of the total variance in behavior among the regional participants (*R*^2^ = 0.142, *p* < 0.001).

## Discussion

In the present study, we surveyed genomic literacy among large numbers of participants from regional and national samples to expand on previous studies with small sample sizes and to compare our baseline regional data against a national dataset. This study is unique in that it examined attitudes, knowledge, concerns, and behaviors concurrently, with a sole focus on these constructs as they relate to genomics. To our knowledge, no previous work has investigated the unique and additive roles of genomic knowledge and attitudes in predicting health-related behaviors.

### Connecting knowledge with application to health

Genomic literacy—understanding enough genomics to be able to communicate effectively and make personal health care decisions—is vital for the public to take advantage of the recent innovations in genomics and its uses in medicine. Our results indicate that the public has some accurate knowledge of basic genetics, answering, on average, 9 of 11 true/false questions correctly. However, participants’ perceived knowledge was lower, and they felt least confident in their understanding of the human genome. This suggests a disconnect between basic factual knowledge and its meaningful application and highlights a key gap in science communication. We found that health care professionals were one of the most utilized sources of information for both medical and genetic research, underscoring a need to make accessible educational materials available to these providers. However, participants were more likely to speak with family members than a physician about genetic testing. This may reflect a distinction in health care professionals, i.e. physician versus other health care professionals such as nurses, occupational therapists etc. Alternatively, there may be a hesitancy to address testing specifically in a health care context. This is worth exploring in the future. Participants also heavily relied upon the internet to learn about genetic research but also turned to magazine and newspaper articles and television. Therefore, there are multiple potential avenues for educating the public.

Attitudes regarding genomic research and its potential to improve health care were overwhelmingly positive, consistent with past results [18]. This was despite a substantial proportion (12–15%) reporting a lack of trust in research agencies and concerns about the consequences, both personal and societal, some of which were reported previously [1, 26, 21, 18]. The regression analyses indicated that knowledge and positive attitudes were associated with an engagement in genomic health behaviors, which may be regarded as effective genomic literacy, that is, the use of one’s “working knowledge of genomic science” to make informed personal decisions [3]. Perhaps not surprisingly, having a genetic disorder or a gene related to a disease also predicted greater engagement in positive genomic health behaviors.

The association between genomic knowledge and engaging in proactive genomic behaviors supports prior claims that the public’s lack of genomic literacy impacts the integration of genomic research into medical practice (such as genetic testing and genome sequencing) [3]. The association also suggests that a broad intervention to improve genomic knowledge will have to be adaptive in increasing public engagement in behaviors, considering the pace of genomic research and its future use in health care. Our findings indicate that a first step in improving genomic literacy would be to connect what people already know about DNA to the concept of the genome and how genomics is being used in medicine to assess risk for disease and to personalize treatments. It appears that trouble integrating these terms and ideas into one’s knowledge base is a major challenge toward genomic literacy.

We also found that knowledge was no longer a predictor of behavior in the regional sample when the analysis included having a genetic disorder and having a gene related to a disease. This may be related to small differences in the educational levels of the regional participants compared to those in the national sample. Alternatively, other variables, such as physician knowledge, might be a factor. Nonetheless, in both datasets, participants that reported having a genetic disorder were more likely to engage in positive genomic health behaviors, which may be attributable to a greater awareness of genomics and a propensity to be proactive about their health. Interestingly, however, having a genetic disorder or a gene related to a disease did not correlate with knowledge. Therefore, other factors likely influence engagement in positive genomic health behaviors, such as health professionals. Those with genetic disorders are likely to have a different relationship with their providers.[33–35] Indeed, more of the variance in behavior in the national sample was accounted for when the analysis included having a genetic disorder or disease related gene. These findings suggest that interventions to improve genomic knowledge will increase the public’s engagement in positive genomic health behaviors. However, our data indicate that educational materials should focus on making connections between genetics, genomics, and health rather than just providing basic facts without context.

### Trust issues

Participants voiced many concerns that may hamper their willingness to engage in positive genomic health behaviors such as participating in research, which has important implications for both genomic research and the expansion of genomics into health care. There are currently initiatives to include more people from diverse backgrounds in genomic research in an effort to better determine genetic and environmental determinants of disease [36, 37], but these efforts rely on the willingness of people to donate their samples to research. Many of the present study’s participants expressed a distrust of relevant authorities (i.e., government, research institutions, and pharmaceutical companies) and concerns about how their samples would be handled. To address these concerns, there needs to be greater transparency and clear disclosure of how samples will be used, which has been cited previously as a key factor in generating trust [38, 39]. Notably, some knowledge of genetics also predicted the likelihood to participate in genome sharing [40, 38], consistent with our findings.

Of note, the surveys in the present study were conducted in 2017, prior to the COVID-19 pandemic, indicating that the tendency for mistrust, at least with respect to genetics and genomic data, predates the pandemic. This is consistent with several reports and articles published in that time frame [41, 42]. Trust in institutions and scientists has since become more complex and often more polarized [43, 44], but trust in health care workers has generally increased, though levels vary by age, ethnicity, community type, etc. [45]. Institutions and agencies should strive to strengthen relationships and develop partnerships with local communities as a way to build trust, to enhance genomic literacy and to encourage public contribution to genomic and medical research.

Participants also had concerns about potential discrimination by health insurance companies. The fear that a risk for disease could be used against an individual would likely deter many from seeking genetic testing. Efforts should therefore be made to advertise the Genetic Information Nondiscrimination Act (GINA) passed in 2008, which prevents health insurance companies from raising premiums on the basis of genetic information and blocks employers from using this information to discriminate against employees or applicants [26]. If health care providers hope to expand the use of genetic testing to improve human health, this information should be communicated with patients. Providers also need to be transparent with patients about the risks; GINA does not apply to life insurance, disability insurance, and other types of supplemental insurance [46–48].

### Limitations of this study

One limitation to consider in interpreting the results of this study is the demographics of our samples. The samples were skewed toward older participants, likely because the survey was administered via landline telephones. Thus, our unweighted samples may not be representative of US or regional (Erie County, NY) populations. It should also be noted that causal claims cannot be made as a result of the cross-sectional nature of this study.

The measures used in this study contained items used in previous research [29, 30]. Although the behavior and perceived knowledge scales demonstrated good levels of reliability, the knowledge and attitudes scales did not. However, the knowledge scale contained several items from a scale that demonstrated good reliability [27, 28]. It is possible that items unique to this study (i.e. questions concerning GMOs and vaccination) detracted from the score’s internal consistency. Principal component analysis of the answers to knowledge questions for each participant indicated that several of the questions do not correlate with each other (Fig. S1). The true/false nature of these questions, which reduces the variability of the answers, may also contribute to the lower reliability coefficient. The attitude scale used in this study could be improved with additional questions about specific elements of genomic research and genetic testing, such as funding or social justice issues. Thus, there is a need, and a movement in current research, to develop and validate high-quality measures of genomic knowledge, attitudes, and behavior [49, 50]

The originality of some of our measures of knowledge and behaviors may make interpretation more difficult. For example, we surveyed participants about behaviors they already performed as well as their intentions to perform them in the future. Additional investigations could expand on this by using other ways of assessing intentions and behaviors. For example, more specific questions could be developed to collect more in-depth information and perhaps focus on specific populations such as those who have received genetic testing. Different measures of knowledge have been used in studies of genomic literacy, making it difficult to compare results. The Wellcome Trust Survey [32, 18] and Lanie and colleagues [19] asked participants only a few simple questions about genes and DNA. Haga and colleagues [21] administered the most extensive and perhaps most difficult survey of genomic knowledge, asking questions about both scientific facts and disease-related concepts. Our participants may have appeared more knowledgeable because we asked mostly basic questions about genetics, although the questions were similar to those used in [32], which reported significantly lower knowledge levels; only a few questions asked about health implications or interactions between genes and the environment. To better determine what the public knows, further studies and those with a more standardized way of assessing genomic knowledge are needed [19, 21, 22].

### Future directions

Once gaps in the public’s knowledge of genomics are recognized, studies can investigate what kinds of education and outreach interventions effectively improve genomic literacy and whether improved knowledge translates to greater engagement in positive genomic health behaviors. The public’s concerns regarding genomic research and its application in medicine should also be explored more deeply. In 2010, McBride and colleagues [24] asked, “What will individuals and society need in order to develop the genetic literacy sufficient for ensuring balanced consideration of current and anticipated genomic discovery?” This question is still relevant today, and genomic researchers should collaborate with experts in public health and communications to answer this question and prepare the public for the incorporation of genomic research into health care. All in all, future research will need to go beyond identifying what the public knows about genomics to understanding how they learned what they know, what they need to know, their nuanced concerns about the social and health implications of genomic research, and what researchers and health care providers can do to build trust and to improve the public’s genomic literacy.

## Acknowledgments

This work was funded by University at Buffalo’s Genome, Environment and Microbiome Community of Excellence. We are grateful to Dr. Karen Dietz for critical reading of the manuscript and expert editing advice. We thank all the anonymous survey participants who took the time to respond thoughtfully to our questions.

