## Supplemental figure and tables for "Challenges in Improving Genomic Literacy: Results from National and Regional Surveys of Genomic Knowledge, Attitudes, Concerns, and Behaviors"

**Supplementary Materials**

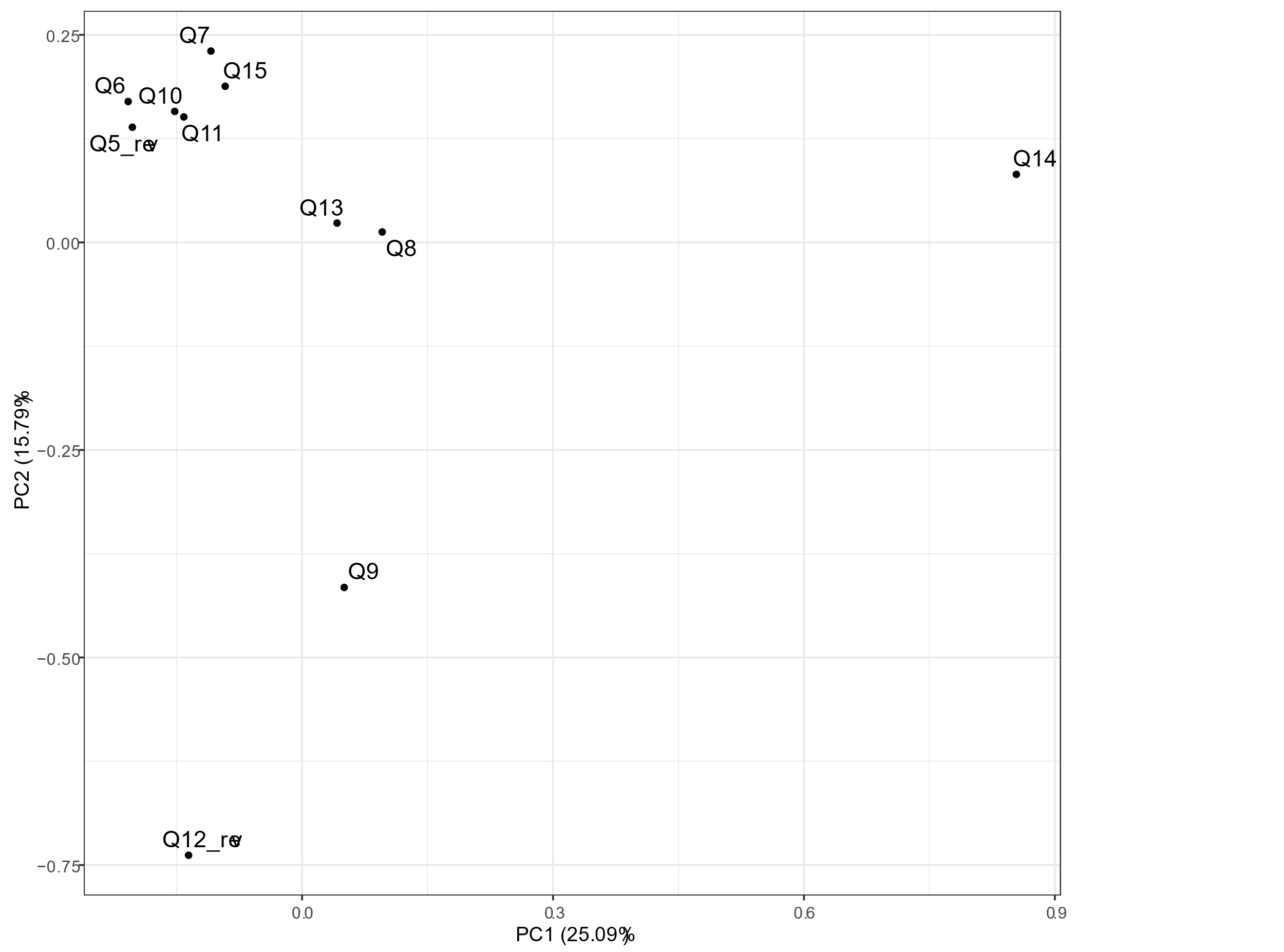

**Figure S1**. Principal components analysis performed on answers of each participant to knowledge questions in **Table S2 (Q.5 to Q.15)**. Some questions cluster, while others segregate away from other questions.

**Table S1.** Perceived knowledge of genomics

| **Term^a^** | **National** | | | **Regional** | |
| --- | --- | --- | --- | --- | --- |
|  | ***n*** | | **%** | ***n*** | **%** |
| Q.1 Human DNA |  |  | |  |  |
| 1 (Have not heard the term) | 39 | 2.6 | | 62 | 6.2 |
| 2 (Little understanding) | 104 | 6.9 | | 81 | 8.1 |
| 3 (Some understanding) | 252 | 16.8 | | 217 | 21.7 |
| 4 (Good understanding) | 438 | 29.2 | | 240 | 24.0 |
| 5 (Excellent understanding) | 667 | 44.5 | | 399 | 39.9 |
| Q. 2 Genetic testing |  |  | |  |  |
| 1 (Have not heard the term) | 75 | 5.0 | | 92 | 9.2 |
| 2 (Little understanding) | 145 | 9.7 | | 93 | 9.3 |
| 3 (Some understanding) | 338 | 22.5 | | 246 | 24.6 |
| 4 (Good understanding) | 445 | 29.6 | | 254 | 25.4 |
| 5 (Excellent understanding) | 498 | 33.2 | | 315 | 31.5 |
| Q.3 Genetically modified organisms (GMOs) |  |  | |  |  |
| 1 (Have not heard the term) | 159 | 10.6 | | 197 | 19.7 |
| 2 (Little understanding) | 151 | 10.1 | | 163 | 16.3 |
| 3 (Some understanding) | 364 | 24.2 | | 225 | 22.5 |
| 4 (Good understanding) | 434 | 28.9 | | 190 | 19.0 |
| 5 (Excellent understanding) | 392 | 26.1 | | 225 | 22.5 |
| Q.4 The human genome |  |  | |  |  |
| 1 (Have not heard the term) | 249 | 16.6 | | 331 | 33.1 |
| 2 (Little understanding) | 232 | 15.5 | | 158 | 15.8 |
| 3 (Some understanding) | 391 | 26.1 | | 235 | 23.5 |
| 4 (Good understanding) | 348 | 23.2 | | 145 | 14.6 |
| 5 (Excellent understanding) | 280 | 18.7 | | 130 | 13.0 |

^a^Understanding of the term was rated on a scale of 1–5.

**Table S2.** Actual knowledge of genomics: correct response frequencies

| **Query^a^** | **Correct response** | | | |
| --- | --- | --- | --- | --- |
|  | **National** | | **Regional** | |
|  | ***n*** | **%** | ***n*** | **%** |
| Q.5 A gene is a disease | 1,472 | 98.0 | 953 | 95.3 |
| Q.6 Our genes control hereditary characteristics, that is, those characteristics inherited from our parents | 1,416 | 94.3 | 950 | 95.0 |
| Q.7 The genes we are born with have the ability to determine heart disease, diabetes, and cancer | 1,334 | 88.8 | 893 | 89.3 |
| Q.8 A gene is a piece of DNA | 1,219 | 81.2 | 780 | 78.0 |
| Q.9 Different body parts have the same genes | 1,085 | 72.3 | 711 | 71.1 |
| Q.10 A person can have a gene for a specific disease and still be completely healthy | 1,395 | 93.0 | 941 | 94.1 |
| Q.11 All plants and animals have DNA | 1,393 | 92.8 | 884 | 88.4 |
| Q.12 Eating genetically modified animal/food products can cause a person’s genes to change | 1,174 | 78.2 | 708 | 70.8 |
| Q.13 The human genome refers to the person’s complete set of DNA | 1,164 | 77.5 | 771 | 77.1 |
| Q.14 Genes can be altered by the environment | 840 | 55.9 | 506 | 50.6 |
| Q.15 The onset of certain diseases is due to a combination of genes, environment, and lifestyle | 1,382 | 92.1 | 873 | 87.3 |

^a^Participants were asked to indicate if the statement was true or false.

**Table S3.** Attitudes regarding genomic research and genomic science

| **Category** | **National** | | | **Regional** | | |
| --- | --- | --- | --- | --- | --- | --- |
|  | ***n*** | | **%** | ***n*** | | **%** |
| Q.16 Do you believe research on the human genome will help improve human health? |  |  | |  |  | |
| Strongly disagree | 13 | 0.9 | | 17 | 1.7 | |
| Disagree | 15 | 1.0 | | 25 | 2.5 | |
| Neutral | 110 | 7.3 | | 90 | 9.0 | |
| Agree | 534 | 35.5 | | 422 | 42.2 | |
| Strongly agree | 830 | 55.3 | | 445 | 44.6 | |
| Q.17 How optimistic are you about the possibility of improved health care as a result of genetic research? |  |  | |  |  | |
| Not at all optimistic | 40 | 2.6 | | 31 | 3.1 | |
| Not too optimistic | 109 | 7.3 | | 52 | 5.2 | |
| Neither | 145 | 9.7 | | 133 | 13.4 | |
| Optimistic | 710 | 47.3 | | 457 | 45.7 | |
| Extremely optimistic | 497 | 33.1 | | 326 | 32.6 | |

**Table S4.** Participants’ concerns regarding genetic research

| **Concern** | **National** | | **Regional** | |
| --- | --- | --- | --- | --- |
|  | ***n*** | **%** | ***n*** | **%** |
| Prompted |  |  |  |  |
| I do not have any concerns about genetic research | 600 | 39.9 | 454 | 45.4 |
| Results could lead to discrimination in my employment | 91 | 6.1 | 60 | 6.0 |
| Results could affect my health insurance premiums | 114 | 7.6 | 99 | 9.9 |
| If I participate in a study, my genetic information might not be kept confidential | 137 | 9.1 | 79 | 7.9 |
| My genetic information may be used for some medical discovery that I may not profit from | 85 | 5.7 | 51 | 5.1 |
| People might use genetic research to create “designer babies” | 220 | 14.6 | 153 | 15.3 |
| Unprompted |  |  |  |  |
| I do not trust authorities in medicine, pharmaceutical companies, and/or government | 239 | 15.9 | 95 | 9.5 |
| More research needs to be done and financially supported by the government | 104 | 6.9 | 58 | 5.8 |
| Concerned about creation of new life and diseases, cloning, modified genes, and organisms | 96 | 6.4 | 66 | 6.6 |
| I do not believe genetic research, genetic testing, and/or use of personal genetic information will be properly regulated or made transparent | 96 | 6.4 | 51 | 5.1 |
| Concerned that information will be used for genetic-based discriminative actions such as eugenics | 67 | 4.5 | 40 | 4.0 |
| Concerned about other forms of discrimination not already mentioned | 66 | 4.4 | 20 | 2.0 |
| Concerns based on conflict with personal religious beliefs | 58 | 3.9 | 30 | 3.0 |
| Concerned about agricultural GMOs | 33 | 2.2 | 8 | 0.8 |
| Concerned about nonconsensual physical human experimentation | 20 | 1.3 | 12 | 1.2 |
| Others | 146 | 9.7 | 49 | 4.9 |

**Table S5.** Participants’ concerns regarding genetic testing and use of personal genetic information

| **Concern** | **National** | | **Regional** | | |
| --- | --- | --- | --- | --- | --- |
|  | ***n*** | **%** | ***n*** | | **%** |
| Prompted |  |  |  |  | |
| I do not have any concerns about genetic testing | 613 | 40.8 | 418 | 41.8 | |
| I don’t have any concerns about the potential use of my genetic information | 530 | 35.3 | 418 | 41.8 | |
| I am concerned that wealthier people could benefit more from genetic testing | 103 | 6.9 | 66 | 6.6 | |
| I am uncomfortable giving samples for genetic testing | 99 | 6.6 | 63 | 6.3 | |
| I am uncomfortable giving samples for genetic research | 97 | 6.5 | 63 | 6.3 | |
| I am concerned about what happens with my samples after the testing is completed | 271 | 18.0 | 195 | 19.5 | |
| I am concerned that researchers will find that I have a gene for a disease | 60 | 4.0 | 53 | 5.3 | |
| I am concerned that I might be predisposed to a disease for which there is no treatment or cure | 65 | 4.3 | 54 | 5.4 | |
| Unprompted |  |  |  |  | |
| I do not trust authorities in medicine, pharmaceutical companies, and/or government | 215 | 14.3 | 133 | 13.3 | |
| I do not believe genetic research, genetic testing, and/or use of personal genetic information will be properly regulated or made transparent | 197 | 13.1 | 101 | 10.1 | |
| Concerned that information will be used for genetic-based discriminative actions such as eugenics | 39 | 2.6 | 19 | 1.9 | |
| Concerned about other forms of discrimination or bigotry not already mentioned | 182 | 12.1 | 78 | 7.8 | |
| More research needs to be done and financially supported by the government | 17 | 1.1 | 4 | 0.4 | |
| Concerned about creation of new life and diseases, cloning, modified genes, and organisms | 17 | 1.1 | 16 | 1.6 | |
| Concerns based on conflict with personal religious beliefs | 11 | 0.7 | 2 | 0.2 | |
| Concerned about agricultural GMOs | 4 | 0.3 | 0 | 0.0 | |
| Concerned about nonconsensual human experimentation | 3 | 0.2 | 0 | 0.0 | |
| Others | 148 | 9.8 | 74 | 7.4 | |

**Table S6.** Sources of information

| **Source^a^** | **National** | | | **Regional** | |
| --- | --- | --- | --- | --- | --- |
|  | ***n*** | | **%** | ***n*** | **%** |
| Medical research |  |  | |  |  |
| Television | 657 | 43.7 | | 443 | 44.3 |
| Internet | 792 | 52.7 | | 431 | 43.1 |
| Medical or scientific journal | 601 | 40.0 | | 301 | 30.1 |
| Magazine article | 685 | 45.5 | | 373 | 37.3 |
| Newspaper article | 655 | 43.6 | | 385 | 38.5 |
| Health professional (doctors/nurses) | 780 | 51.9 | | 457 | 45.7 |
| Radio | 304 | 20.2 | | 172 | 17.2 |
| Directly from an advertisement about genetic testing | 225 | 15.0 | | 132 | 13.2 |
| Personal experience | 335 | 22.3 | | 203 | 20.3 |
| Family or friend | 505 | 33.6 | | 310 | 31.0 |
| Other | 95 | 6.3 | | 99 | 9.9 |
| Human genome and/or genetic research |  |  | |  |  |
| Television | 577 | 38.4 | | 344 | 34.4 |
| Internet | 684 | 45.5 | | 373 | 37.3 |
| Medical or scientific journal | 544 | 36.2 | | 236 | 23.6 |
| Magazine article | 583 | 38.8 | | 326 | 32.6 |
| Newspaper articl | 525 | 34.9 | | 333 | 33.3 |
| Health professional (doctors/nurses) | 529 | 35.2 | | 322 | 32.2 |
| Radio | 201 | 13.4 | | 113 | 11.3 |
| Directly from an advertisement about genetic testing | 146 | 9.7 | | 97 | 9.7 |
| Personal experience | 225 | 15.0 | | 142 | 14.2 |
| Family or friend | 350 | 23.3 | | 217 | 21.7 |
| Other | 128 | 8.5 | | 148 | 14.8 |

^a^Participants were asked to select all that apply for their source medical research or human genome and/or genetic research.

**Table S7.** Positive genomic health behaviors

| **Behavior^a^** | **National** | | **Regional** | |
| --- | --- | --- | --- | --- |
|  | ***n*** | **%** | ***n*** | **%** |
| Getting a genetic test |  |  |  |  |
| (1) No, never considered | 790 | 52.6 | 656 | 65.6 |
| (2) Thought about it but decided not to | 155 | 10.3 | 116 | 11.6 |
| (3) Thought about it and plan to in the future | 318 | 21.2 | 90 | 9.0 |
| (4) Yes, already done it | 222 | 14.8 | 129 | 12.9 |
| Missing or declined to answer | 15 | 1.0 | 9 | 0.9 |
| Seeking more information about genetic testing |  |  |  |  |
| (1) No, never considered | 744 | 49.6 | 630 | 63.0 |
| (2) Thought about it but decided not to | 143 | 9.5 | 89 | 8.9 |
| (3) Thought about it and plan to in the future | 308 | 20.5 | 121 | 12.1 |
| (4) Yes, already done it | 284 | 18.9 | 151 | 15.1 |
| Missing or declined to answer | 22 | 1.4 | 9 | 0.9 |
| Speaking to my physician about genetic testing |  |  |  |  |
| (1) No, never considered | 1,015 | 67.6 | 737 | 73.8 |
| (2) Thought about it but decided not to | 79 | 5.3 | 47 | 4.7 |
| (3) Thought about it and plan to in the future | 147 | 9.8 | 64 | 6.4 |
| (4) Yes, already done it | 247 | 16.5 | 142 | 14.2 |
| Missing or declined to answer | 13 | 0.8 | 9 | 0.9 |
| Speaking to a family member or friend about genetic testing |  |  |  |  |
| (1) No, never considered | 735 | 49.0 | 581 | 58.1 |
| (2) Thought about it but decided not to | 70 | 4.7 | 49 | 4.9 |
| (3) Thought about it and plan to in the future | 108 | 7.2 | 69 | 6.9 |
| (4) Yes, already done it | 579 | 38.5 | 295 | 29.5 |
| Missing or declined to answer | 9 | 0.6 | 6 | 0.6 |
| Reading labels to check if a food is genetically modified^b^ |  |  |  |  |
| (1) No, never considered | 620 | 41.3 | 447 | 44.7 |
| (2) Thought about it but decided not to | 59 | 3.9 | 31 | 3.0 |
| (3) Thought about it and plan to in the future | 51 | 3.4 | 38 | 3.8 |
| (4) Yes, already done it | 757 | 50.4 | 475 | 47.5 |
| Missing or declined to answer | 14 | 0.9 | 10 | 1.0 |
| Avoiding recommended vaccinations^b^ |  |  |  |  |
| (1) No, never considered | 1,138 | 75.8 | 791 | 79.2 |
| (2) Thought about it but decided not to | 39 | 2.6 | 23 | 2.3 |
| (3) Thought about it and plan to in the future | 49 | 3.2 | 14 | 1.4 |
| (4) Yes, already done it | 241 | 16.0 | 165 | 16.5 |
| Missing or declined to answer | 34 | 2.3 | 6 | 0.6 |
| Modifying my lifestyle based on a family history of a certain disease |  |  |  |  |
| (1) No, never considered | 423 | 28.2 | 324 | 32.4 |
| (2) Thought about it but decided not to | 74 | 5.0 | 38 | 3.8 |
| (3) Thought about it and plan to in the future | 148 | 9.8 | 66 | 6.6 |
| (4) Yes, already done it | 845 | 56.3 | 562 | 56.2 |
| Missing or declined to answer | 12 | .8 | 10 | 1.0 |
| Participating in genetic research |  |  |  |  |
| (1) No, never considered | 1,041 | 69.3 | 736 | 73.7 |
| (2) Thought about it but decided not to | 86 | 5.7 | 51 | 5.1 |
| (3) Thought about it and plan to in the future | 197 | 13.1 | 82 | 8.2 |
| (4) Yes, already done it | 157 | 10.5 | 124 | 12.4 |
| Missing or declined to answer | 20 | 1.3 | 6 | 0.6 |

^a^Rating of actions/behaviors related to genetic testing on a scale of 1–5.

^b^Not included in the final behavior scale.

**Table S8.** Correlations among demographics, knowledge, attitudes, and behavior in regional dataset

| **Category** | ***r* value** | | | | | | | | |
| --- | --- | --- | --- | --- | --- | --- | --- | --- | --- |
|  | **Gender** | **Education** | **Income** | **Age** | **Knowledge** | **Attitudes** | **Behavior** | **Health** | **Genetic disorder** |
| Gender | – |  |  |  |  |  |  |  |  |
| Education | −0.10** | – |  |  |  |  |  |  |  |
| Income | −0.05 | 0.24*** | – |  |  |  |  |  |  |
| Age | −0.23*** | 0.03 | −0.09** | – |  |  |  |  |  |
| Knowledge | −0.09** | 0.27*** | 0.02 | −0.04 | – |  |  |  |  |
| Attitudes | −0.09** | 0.11*** | 0.04 | 0.06 | 0.30*** | – |  |  |  |
| Behavior | −0.11*** | 0.15*** | 0.05 | −0.01 | 0.20*** | 0.23*** | – |  |  |
| Health | −0.07* | 0.23*** | 0.19** | −0.05 | 0.09** | 0.03 | 0.07* | – |  |
| Genetic disorder | −0.04 | −0.04 | −0.05 | 0.07 | 0.02 | 0.01 | 0.15*** | −0.20*** | – |
| Gene related to a disease | −0.04 | 0.02 | −0.07 | 0.02 | 0.06 | 0.01 | 0.19*** | −0.14*** | 0.44*** |

**p* < 0.05 ***p* < 0.01, ****p* < 0.001.

**Table S9.** Results of hierarchical linear regression using a regional dataset

| **Category** | **β value** | |
| --- | --- | --- |
|  | **Model 1** | **Model 2** |
| Gender | −1.080** | −0.851** |
| Education | 0.547*** | 0.367** |
| Age | −0.111*** | −0.098*** |
| Actual knowledge |  | 0.360** |
| Attitudes |  | 0.563*** |
| Total *R*^2^ | 0.034 | 0.034 |
| Change in *R*^2^ | 0.086*** | 0.052*** |

***p* < 0.01, ****p* < 0.001.

**Table S10.** Results of multiple regression with health questions using a regional dataset

| **Category** | **β value** |
| --- | --- |
| Gender | −1.091** |
| Education | 0.483** |
| Age group | −0.111 |
| Quality of health | 0.125 |
| Genetic disorder | 1.114* |
| Gene related to a disease | 1.843*** |
| Actual knowledge | 0.245 |
| Attitudes | 0.580*** |
| Total *R*^2^ | 0.142*** |

**p* < 0.05, ***p* < 0.01, ****p* < 0.001.
